# Imipridones inhibit tumor growth and improve survival in an orthotopic liver metastasis mouse model of human uveal melanoma

**DOI:** 10.1101/2024.01.12.575058

**Authors:** Chandrani Chattopadhyay, Jason Roszik, Rajat Bhattacharya, Md Alauddin, Iqbal Mahmud, Sirisha Yadugiri, Mir Mustafa Ali, Fatima S. Khan, Varun Vijay Prabhu, Phillip Lorenzi, Elizabeth Burton, Rohini R. Morey, Rossana Lazcano, Michael A. Davies, Sapna P. Patel, Elizabeth A. Grimm

## Abstract

**Purpose:** Uveal melanoma (UM) is a highly aggressive disease with very few treatment options. We previously demonstrated that mUM is characterized by high oxidative phosphorylation (OXPHOS). Here we tested the anti-tumor, signaling and metabolic effects of imipridones, CLPP activators which reduce OXPHOS indirectly and have demonstrated safety in patients.

**Experimental Design:** We assessed CLPP expression in UM patient samples. We tested the effects of imipridones (ONC201, ONC212) on the growth, survival, signaling and metabolism of UM cell lines *in vitro,* and for therapeutic effects *in vivo* in UM liver metastasis models.

**Results:** CLPP expression was confirmed in primary and mUM patient samples. ONC201/212 treatment of UM cell lines *in vitro* decreased OXPHOS effectors, inhibited cell growth and migration, and induced apoptosis. ONC212 increased metabolic stress and apoptotic pathways, inhibited amino acid metabolism, and induced cell death-related lipids. ONC212 also decreased tumor burden and increased survival *in vivo* in two UM liver metastasis models.

**Conclusion:** Imipridones are a promising strategy for further testing and development in mUM.

## Introduction

Uveal melanoma (UM), a rare sub-type of melanoma, is the most common primary cancer of the eye in adults and is diagnosed in about 2,500 adults/year in the United States. About 50% of the UM patients progress to metastatic disease and 90% of these patients show preferential liver metastasis (1–4). UM patients with advanced liver metastasis have poor prognosis with median survival of less than a year (2). Metastatic uveal melanoma (mUM) does not respond to conventional chemotherapy and only rarely responds to immune therapies that are approved for cutaneous melanoma patients (5). mUM patients currently have only two FDA approved therapeutic options: Tebentafusp (which is restricted to HLA-A*02:01-positive adult patients) for patients with unresectable disease (6) and melphalan for liver-directed therapy (7). There remains a need for new therapeutics that can achieve durable responses in mUM patients.

UM is a unique disease and distinct from cutaneous melanoma due to low mutation load and high preference for liver metastasis (>90%) (8). Loss of BRCA-Associated protein-1 (*BAP1*) (9,10) and loss of a copy of chromosome 3 (11) are significant prognostic factors for UM metastasis. G-protein coupled receptors GNAQ and GNA11 are frequently mutated in UM (12,13), but currently approved therapies to target these oncogenes are not available. IGF-1 pathway inhibition has shown to control UM growth in preclinical studies (14), but did not show promising outcome in a clinical study (15). Thus, it is essential to identify novel vulnerabilities downstream of these aberrations to develop effective therapies.

We recently showed that UM has high OXPHOS and high expression of OXPHOS effectors like SDHA in tumors with monosomy 3 UM (M3UM), a finding independently validated by other groups (16,17). Unfortunately, direct OXPHOS inhibition was associated with unacceptable toxicity in patients (18–20) and currently potent inhibitors of OXPHOS effectors are not clinically available. Imipridones are a distinct class of small molecule anti-cancer compounds (21). Their primary mechanism of action is to activate mitochondrial protease CLPP causing dysregulated proteolysis of mitochondrial OXPHOS effectors like SDHA, SDHB, NDUFA12, thereby reducing their cellular levels (22–24). The imipridone compound ONC201 has shown efficacy in several preclinical models of solid tumors and hematologic malignancies, and showed promising safety, pharmacokinetic and pharmacodynamic efficacy profiles in phase I/II trials involving more than 100 patients (22,25). ONC212, a more potent analog of ONC201, has been evaluated in multiple preclinical studies including in Acute Myelodysplastic Leukemia (AML) (26,27). The anti-cancer activity of imipridones demonstrated in preclinical studies, along with promising early clinical data, suggest that these compounds may present an important new agent for cancer therapy in the future. Their development will be strengthened by an improved understanding of their activity in different cancer types.

In this study, we tested ONC201 and ONC212 for their ability to modulate OXPHOS effectors in UM *in vitro* and *in vivo*. Our studies show that *in vitro* treatment with ONC201 and ONC212 decreased OXPHOS effector proteins, inhibited UM cell growth, induced apoptosis, and inhibited cell migration. ONC212, which was more potent than ONC201 *in vitro,* significantly upregulated mTOR and apoptotic pathways, altered protein and redox metabolism, and increased cell death associated lipids. Importantly, ONC212 inhibited tumor growth and improved survival *in vivo* in orthotopic UM liver metastasis mouse models. Together, our data suggest that ONC212 may be a promising agent for clinical development in UM.

## Materials and Methods

### Antibodies and Reagents

The antibodies for caspase 3 (RRID: AB_2827742), SDHA (RRID:AB_301433), and CLPP (RRID:AB_10975619) were from Abcam, caspase 9 (RRID:AB_2068621), cleaved PARP (RRID:AB_10699459) and β1 integrin (RRID: AB_823544 ) were from Cell signaling Technology, β1-actin (RRID:AB_626632) and SDHB (RRID:AB_10659104) were from Santa Cruz Biotechnology, and F-Actin from Bioss Inc., (RRID:AB_10859354).

ONC201 and ONC212 were provided by Oncoceutics/Chimerix under an MTA.

Cell migration assay plates were from Corning (354578; Corning) and the staining kit was from Siemens Health care Diagnostic Inc. (B4132-1A). Methylthiazole tetrazolium (MTT) reagent was from EMD Millipore Corp (475989), Propidium iodide (81845) and ribonuclease A (RNase; R4642) from Sigma Aldrich.

### Cell Culture and Treatments

Cell line MEL20–06–039, (RRID:CVCL_8473) (28,29), was obtained from Dr. Tara A. McCannel. Cell lines OMM-1 (RRID:CVCL_6939) (30), MEL202 (RRID: CVCL_C301), 92.1 (RRID: CVCL_8607) (30), and MEL270 (RRID: CVCL_C302) (30) were kindly provided by Drs. Martine Jager and Bruce Ksander. UM cell lines obtained from American Type Culture Collection (ATCC) were: MM28 (#CRL-3295), MP38 (#CRL-3296), MP41 (#CRL-3297), MP46 (#CRL-3298), MP65 (#CRL-3299). Additional UM cell line information is provided in Figure S1A (31–34). UM cells were cultured in RPMI 1640 media with 10% FBS, 1% glutamine, 1% Penicillin-streptomycin, and 1% Insulin supplement, under ambient oxygen at 37 °C. Cell lines were validated by short random repeat (STR) DNA fingerprinting techniques and mutational analysis, by the MDACC Cancer Center Support Grant (CCSG)-supported Characterized Cell Line Core.

### Western Blotting

Cells were lysed in a buffer containing 50 mM Tris (pH 7.9), 150 mM NaCl, 1% NP40, 1 mM EDTA, 10% glycerol, 1 mM sodium vanadate, and a protease inhibitor cocktail (Roche Pharmaceuticals). Proteins were separated by SDS-PAGE (Bio-Rad Laboratories), transferred to a Hybond-ECL nitrocellulose membrane (GE Healthcare Biosciences) and blocked in 5% milk. After primary and secondary antibody incubation Thermo Scientific Super Signal chemiluminescence reagent was used for detection, and Li-COR C-digit Scanner was used for imaging and quantitation. For each marker, the western blots were standardized multiple times and appropriate representative blots are included in the figures.

### Cell Cycle Analysis Using Propidium Iodide Staining and FACS

UM cells were trypsinized, washed, and fixed with 70% Ethanol and kept at 4 °C overnight. Cells were centrifuged and pellets were resuspended in PBS to rehydrate for 15 min. Cells were then centrifuged at 500× *g* and treated with 200 µg/mL RNase A for 1 h at 37 °C. Cells were stained with 40 µg/mL Propidium Iodide for 20 min at room temperature. After centrifugation at 500× *g*, cells were resuspended in PBS with 0.02% EDTA for cell cycle analysis using a Beckman Coulter Galios 561 analyzer (Beckman Coulter).

### Colony Formation Assay

For the colony formation assay, UM cells were seeded in 24 well plates at 500 cells/well, and ONC212 (0.1, 0.25 and 0.5 µM) was added the following day. Controls were untreated cells. The drug was replaced every three days. Colonies were grown until control wells reached 70–80% confluency. Cells were washed with 1X PBS and stained with 1 mL of crystal violet in 25% methanol for 5 min at room temperature. Colonies were imaged and counted. The effect of ONC212 on colony formation was quantitated by counting colonies from 10 independent fields/sample and was plotted as a bar graph.

### Reverse Phase Protein Array (RPPA) Analysis

RPPA analyses were performed at The University of Texas MD Anderson Cancer Center Functional Proteomics RPPA Core facility(14). Briefly, cell lysates were serially diluted (from undiluted to 1:16 dilution) and arrayed on nitrocellulose-coated slides. Samples were probed with antibodies using catalyzed signal amplification and visualized by 3,3’-diaminobenzidine colorimetric reaction. Slides were scanned on a flatbed scanner to produce 16-bit TIFF images and were quantified using the MicroVigene software program (Version 3.0). Relative protein levels for each sample were determined by interpolation of each dilution curve. Heatmaps were generated in Cluster 3.0 (http://www.eisenlab.org/6isen/ (accessed on 08/11/2023) as a hierarchical cluster using Pearson correlation and a center metric. RPPA data were normalized by antibody then by sample.

For pathway analysis, RPPA data were median-centered and normalized by standard deviation across all samples for each component to obtain the relative protein level. The pathway score is then the sum of the relative protein level multiplied by its weight of all components in a particular pathway (35).

### Cell Viability Assays

MTT- based cell viability assays were used for estimating cell survival. UM cells were plated at a density of 1 × 10^4^ cells/well in triplicate in a 24-well plate. Next day, ONC201 and/or ONC212 was added to the cells in designated doses and incubated for 72 h. To assess cell viability, MTT reagent dissolved in PBS was added to wells for a final concentration of 1 mg/mL. After 2 h the precipitate formed was dissolved in DMSO, and the color intensity was estimated in an MRX Revelation microplate absorbance reader (Dynex Technologies) at 570 nm.

### Cell Migration Assay

Cell migration assays were performed in Boyden chambers using uncoated filters (BD Biocoat control inserts). UM cells were plated and treated overnight with ONC212 (0.2 µM). Untreated cells were used as a control. The next day, 1 × 10^5^ cells/well were plated in a serum-free medium, and the migration assay was completed with 10% FBS as a chemo-attractant and procedure as described in Chattopadhyay et al. (36). Stained cells were photographed with a Nikon Eclipse TE2000-U microscope (Nikon Instruments Inc.) at 20x magnification using Nikon’s NIS Elements advanced research software. To quantify migration, the cells in each filter were counted from five independent fields under the microscope at 40x magnification and the mean cell number/field was calculated. Each assay condition was tested in duplicates.

### Metabolomics Profiling

Metabolites were extracted using 1 mL of ice-cold 0.1% ammonium hydroxide in 80/20 (v/v) methanol/water. Extracts were centrifuged at 17,000 g for 5 min at 4 °C , the evaporated supernatants were reconstituted in deionized water, and 10 μL were injected for analysis by ion chromatography – mass spectrometry (IC-MS). Data were acquired using a Thermo Orbitrap Fusion Tribrid Mass Spectrometer under ESI negative ionization mode at a resolution of 240,000. Raw data files were imported to Thermo Trace Finder software for final analysis. The relative abundance of each metabolite was normalized by total area, log transformed, and scaled by z-score. Metabolite data were processed and annotated using Thermo Scientific Compound discoverer software (version 3.3 SP2) and relative abundance and metabolic pathway analyzed using R scripts written in house.

### Lipidomics Profiling

To each cell sample, 200 µL of extraction solution containing 2% Avanti SPLASH® LIPIDOMIX® Mass Spec Standard, 1% 10 mM butylated hydroxytoluene in ethanol was added and vortexed 10 min, incubated on ice for 10 min and centrifuged at 13,300 rpm for 10 min at 4 °C. 10 µL of supernatant was injected. Mobile phase A (MPA) was 40:60 acetonitrile: water with 0.1% formic acid and 10 mM ammonium formate. Mobile phase B (MPB) was 90:9:1 isopropanol: acetonitrile: water with 0.1% formic acid and 10 mM ammonium formate. The chromatographic method included a Thermo Fisher Scientific Accucore C30 column (2.6 µm, 150 x 2.1 mm) maintained at 40 °C, autosampler tray chilling at 8 °C, a mobile phase flow rate of 0.200 mL/min, and a gradient elution program as follows: 0-3 min, 30% MPB; 3-13 min, 30-43% MPB; 13.1-33 min, 50-70% MPB; 48-55 min, 99% MPB; 55.1-60 min, 30% MPB. A Thermo Fisher Scientific Orbitrap Fusion Lumos Tribrid mass spectrometer was operated in data dependent acquisition mode, in both positive and negative ionization modes, with scan ranges of 150 – 827 and 825 – 1500 m/z. An Orbitrap resolution of 120,000 (FWHM) was used for MS1 acquisition and a spray voltage of 3,600 and -2900 V were used for positive and negative ionization modes, respectively. Vaporizer and ion transfer tube temperatures were set at 275 and 300 °C, respectively. The sheath, auxiliary and sweep gas pressures were 35, 10, and 0 (arbitrary units), respectively. For MS2 and MS3 fragmentation a hybridized HCD/CID approach was used. Lipid data were processed and annotated using Thermo Scientific LipidSearch software (version 5.0) and analyzed using R scripts written in house.

### Tumor Models for UM Metastasis

#### Generation of Luciferase Tagged UM Cells

The four UM models that were used for generating orthotopic liver growth of UM to determine the effect of ONC212 on tumor growth were tagged with luciferase using lentiviral infections. The lentivirus was purchased from GenTarget Inc. (LVP020-PBS).

#### Splenic Injection

We followed the splenic injection methodology as described in Sugase et al (37). Briefly, NSG mice (RRID:IMSR_JAX:005557; from Jackson Laboratory, Bar Harbor, ME)were placed in the right lateral recumbent position. A 1-cm incision was made in the left upper abdominal wall, followed by a 1 cm incision in the peritoneum to expose the spleen. 0.5 × 10^6^ cells in 50 μL of HBSS were gently injected into the spleen. The insertion site of the needle was cauterized and sealed with sutures to curtail bleeding. Splenectomy was performed 15 min after injection using surgical cautery tip (#231; McKesson). The abdominal incision was closed in two layers with 5-0 polydioxanone absorbable thread (AD Surgical). Mice were injected with Buprenorphine SR subcutaneously for pain relief (1 mg/kg) and kept under observation for 72 h.

#### Monitoring Tumor Growth via *In Vivo* Imaging

For luciferin-based imaging, mice were anesthetized by the XGI-8 Gas Anesthesia system (2% isoflurane; Xenogen) (38). After 10 min of intraperitoneal injection of luciferin (150 mg/kg body weight), bioluminescence intensity was measured by In Vivo Imaging System (IVIS)-200 (Xenogen). Image sequence was acquired using Living Image (Xenogen) software. The images were cropped to fit the final figures using Adobe Photoshop Software.

#### ONC212 Treatment

After confirming tumor growth in the liver, mice were randomized for treatment with ONC212. ONC212 was dissolved in sterile water fresh on the day of treatment and the mice were treated with 25 mg/kg ONC212 twice weekly via oral gavage. The mice in the vehicle group were treated with sterile water.

#### Immunohistochemistry

Formalin-fixed, paraffin-embedded samples were sectioned at 4 micron thickness onto Superfrost plus slides (Fisher Scientific). Slides were stained with the Bond Polymer Refine Red Detection kit using a Bond Rxm automated stainer (Leica Biosystems). Following a 20 min antigen retrieval in pH 6 citrate buffer (CLPP) or pH 9 Tris-EDTA buffer (SDHA) and peroxidase block in 3% hydrogen peroxide, slides were incubated with diluted primary antibody. CLPP was diluted 1:1,000 in Bond primary antibody diluent (Leica, #AR9352), and SDHA was diluted 1:10,000 in Dako antibody diluent (Agilent #S0809). Detection was performed per the standard protocol for the Bond Refine kit. A pathologist scored the IHC-stained slides, evaluating the cytoplasmic staining in the tumor cells. The results were registered as intensity; 0=negative, 1=low, 2=high, and extent; 0 - 100%, multiplying intensity and extent to derive a score from 0 - 200.

#### Statistical Analysis

Both metabolomics and lipidomics data were first normalized by total area normalization followed by log2 transformation and autoscaling. Hierarchical clustering heatmap was performed using relative abundance of analytes. RPPA protein expressions were similarly analyzed using heatmaps. Expression differences were also analyzed using t-tests and visualized using the Tableau Desktop software (v8.1). Tumor growth figures were created with GrapPad Prism (v9). Survival analyses were performed and log-rank p-values were calculated in R (v3.6.0) using the ’survival’ package.

#### Data Availability Statement

All relevant data are available from the corresponding author upon request.

## Results

### CLPP is expressed in primary and metastatic UM

Mitochondrial CLPP is the known target of imipridones (21,22). Thus, we investigated the expression of CLPP in UM. CLPP gene expression was analyzed in 31 unique tumor types and their corresponding normal tissues, including primary UM tumors from The Cancer Genome Atlas (TCGA). UM tumors showed high CLPP gene expression amongst the evaluated cancer types (Figure 1A). Further analysis of the TCGA UM data set revealed higher expression of CLPP in monosomy 3 (M3) compared to disomy 3 (D3) UM (p <0.0001; Figure 1B).

**Figure 1.**
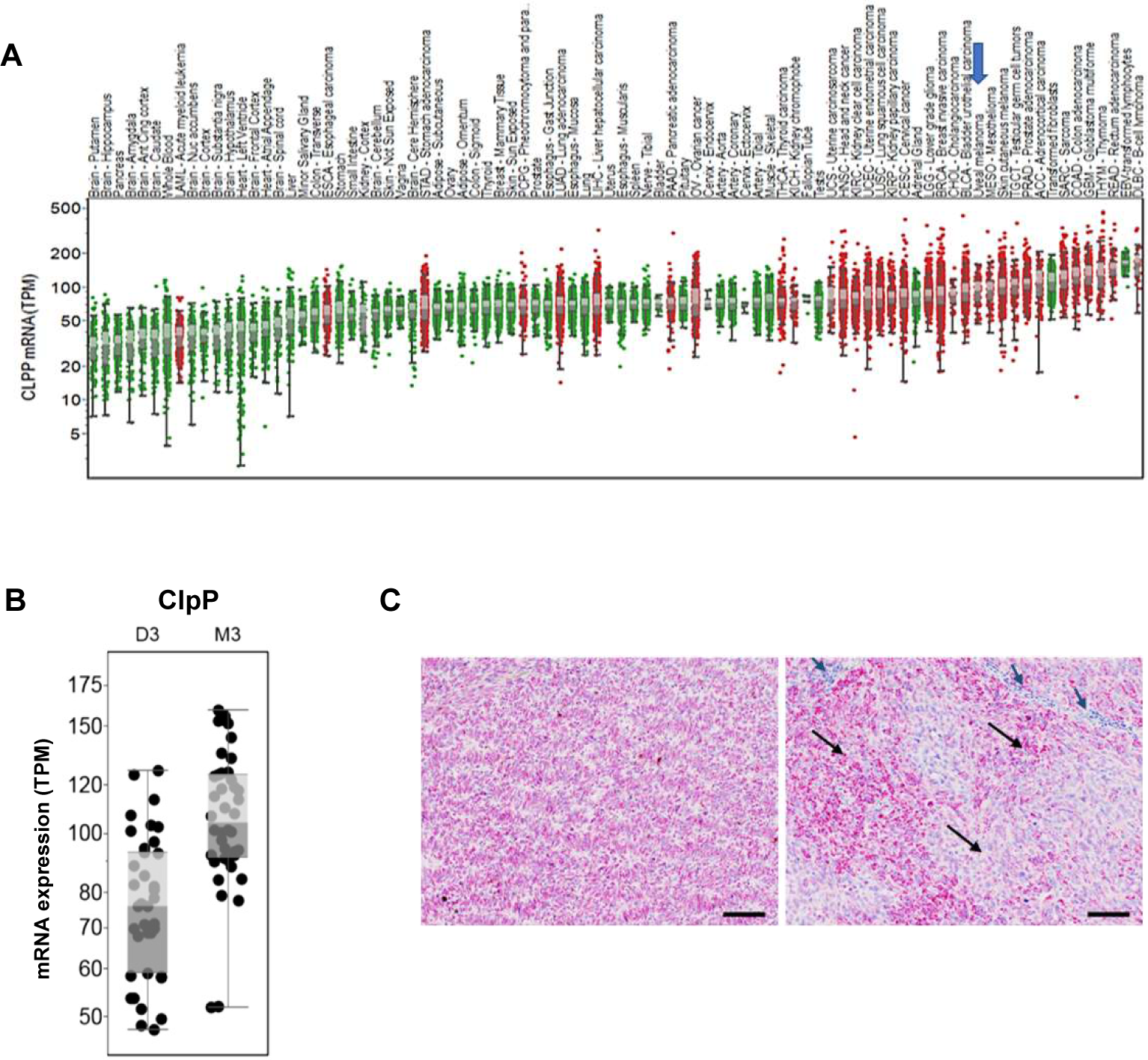
CLPP expression in primary and metastatic UM. (A) The ClpP mRNA expression levels in tumor and normal tissues were compared in 33 cancer (red dots) and corresponding normal (green dots) tissues through TCGA database analysis. The expression in UM is indicated by a blue arrow. **(B)** ClpP mRNA expression of M3 (n = 42) vs. D3 (n = 38) UM tumors from the UM TCGA. Box plots show median values and the 25th to 75th percentile range of the data (i.e., the interquartile range). **(C)** IHC for CLPP protein expression in mUM liver metastases from patients (10X magnification; scale bar = 100 µm). The blue arrows indicate stromal cells and lymphocytes, and the black arrows indicate the regions with melanoma tumors.

Immunohistochemical analysis of mUM clinical samples (n=30) from various sites detected CLPP protein expression in 100% of tumor cells (representative results shown in Figure 1C). High expression (H-score >100) of CLPP was observed in 24 (80%) samples, which included soft tissue, liver, skin, lung, small intestine, brain, and lymph node metastases. Tumor tissues showed a higher intensity of CLPP than adjacent normal tissue in all samples.

### Imipridones reduce mitochondrial OXPHOS effectors and inhibits cell survival in UM cells

Mitochondrial CLPP regulates OXPHOS through proteolysis of effector proteins such as SDHA and SDHB (39). Imipridones indirectly regulate OXPHOS levels through modulation of CLPP activity. Thus, the effect of imipridone treatment on SDHA and SDHB in UM cells was determined. ONC201 and ONC212 reduced the levels of SDHA and SDHB in UM cell lines (MEL20-06-039, MP41, OMM2.3, and 92.1) in vitro by western blotting analysis (Figure 2A – 2B). ONC212 inhibited the expression of SDHA and SDHB more potently than ONC201. As expected, the levels of CLPP expression were not altered with imipridone treatment.

**Figure 2.**
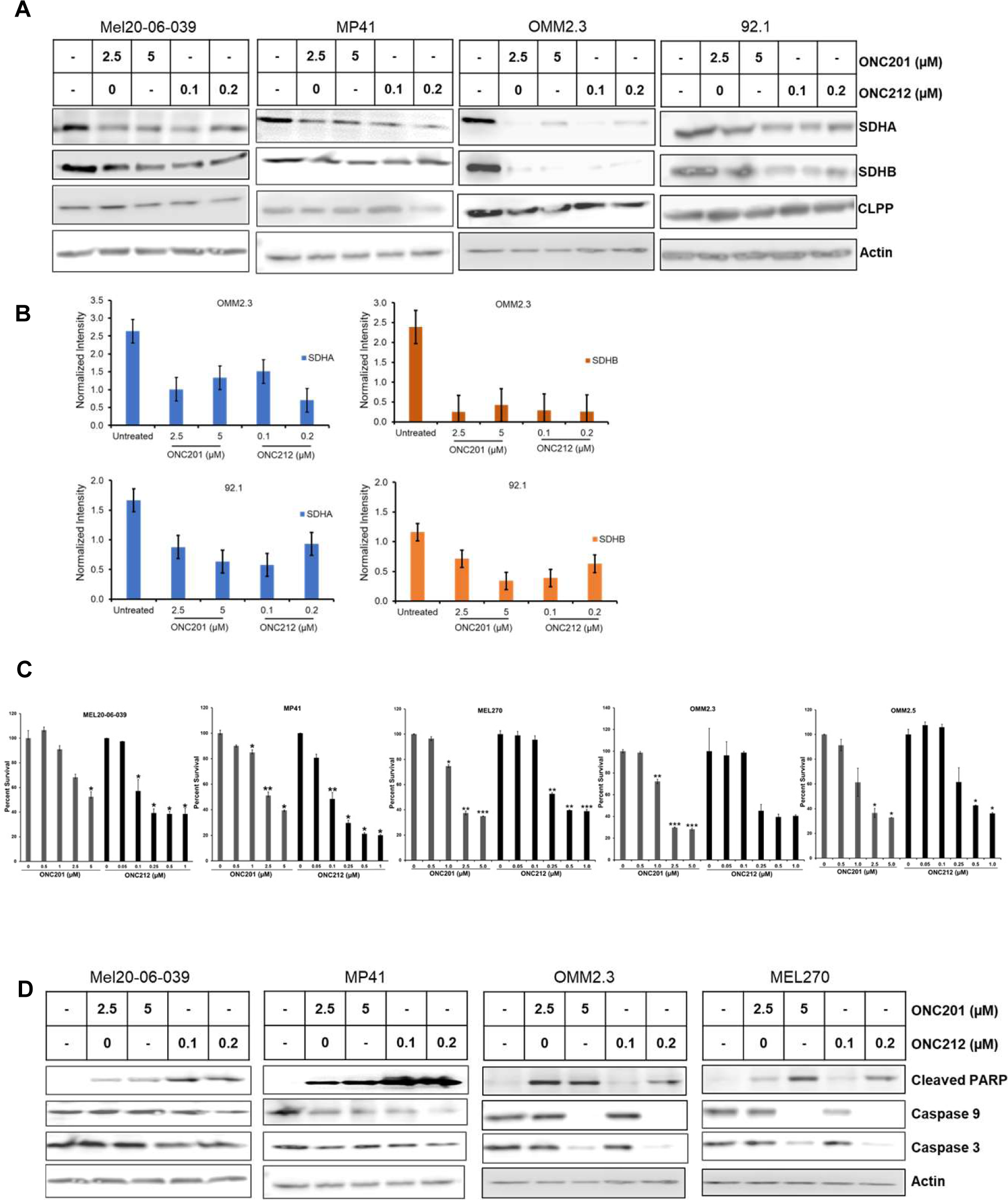
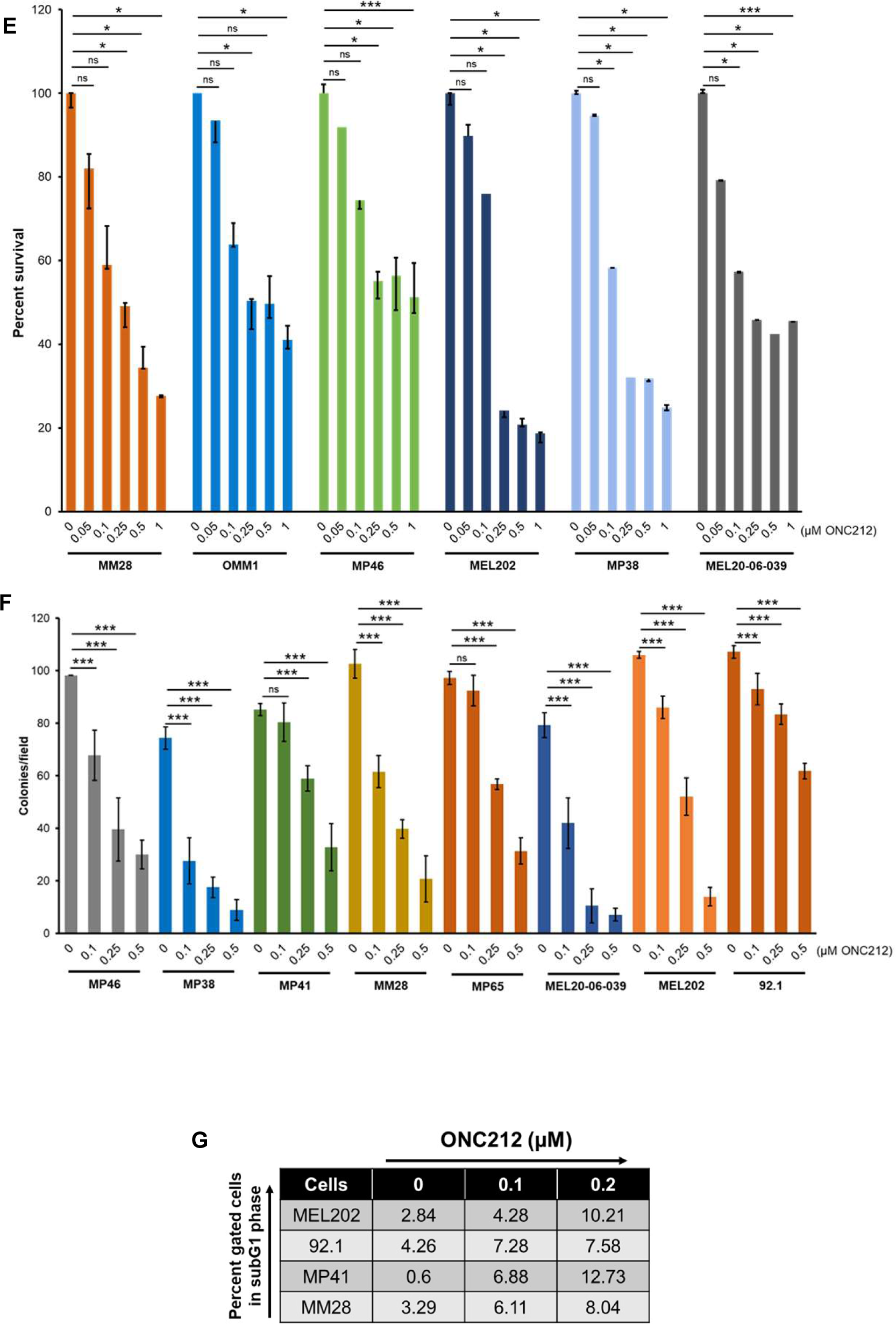
Imipridone treatment reduces OXPHOS effector proteins, inhibits cell survival, and induces apoptosis in UM cell lines. (A) Western blot analysis of UM cell lines (MEL20-06- 039, MP41, OMM2.3, and 92.1) treated with ONC201 or ONC212 for 48 h. **(B)** Quantitation of SDHA and SDHB expression levels from the western blots in 2A. Actin was used for normalization. **(C)** Effect of ONC201 and ONC212 on UM cell survival (in MEl20-06-039, MP41, MEL270, OMM2.3, and OMM2.5 cell lines) by MTT-based cell survival assay. Bar graphs, mean ± SD of three independent experiments. p-values were calculated by comparison to untreated controls (* p < 0.05; ** p < 0.01; *** p < 0.001). **(D)** Western blot analysis of cleaved PARP, intact Caspase 9, and intact Caspase 3 following treatment with ONC201 and ONC212. **(E)** MTT- based measurement of dose-dependent growth inhibition of UM cell lines (MM28, OMM1, MP46, MEL202, MP38, and MEL20-06-039) by ONC212. Bar graphs, mean ± SD of three independent experiments. p-values were calculated by comparison to untreated controls (* p < 0.05; ** p < 0.01; *** p < 0.001). **(F)** Colony formation assay: The average number of colonies observed after staining with crystal violet in ONC212 treated (0.1, 0.25 and 0.5 µM) and untreated control UM cell lines (MP46, MP38, MP41, MM28, MP65, MEL20-06-039, MEL202, and 92.1) were plotted. A total of 5 fields/condition/cell line were counted to obtain the average number of colonies. The bar graphs represent average number of colonies per field and are a mean ± SD of three independent experiments. p-values were calculated by comparing untreated controls with ONC212 treatment doses (* p < 0.05; ** p < 0.01; *** p < 0.001). **(G)** FACS analysis of PI-stained cells collected after 48 h treatment with ONC212.

We next evaluated the impact of ONC201 and ONC 212 on cell proliferation and survival *in vitro*. ONC212 was approximately 10-fold more potent in inhibiting UM cell viability in MTT assays than ONC201 (Figure 2C). Both compounds also induced apoptosis, as demonstrated by increased levels of cleaved PARP (Figure 2D), as well as reduced levels of unprocessed caspase 3 and 9. Similar to cell viability, ONC212 was more potent in inducing apoptosis than ONC201.

Based on these initial results, we focused further experiments on the characterization of ONC212 in UM. ONC212 significantly inhibited UM cell growth in vitro in a dose-dependent manner in all UM cell lines tested (Figure 2E). IC_50_ concentrations ranged from 60 -130 nM (Figure S1B). ONC212 also significantly inhibited colony formation in a clonogenic colony formation assay in MP46, MP38, MP41, MM28, MP65, MEL20-06-039, MEL202, and 92.1 UM cell lines (Figure 2F and S1C). Cell cycle analysis (propidium iodide staining) revealed an increase in G01/G1 cell population in an ONC212 dose-dependent manner (Figure 2G and S1D). All these effects were observed at or near IC_50_ doses. The mechanism of ONC212 action in UM cells was determined by siRNA based CLPP knockdown followed by ONC212 treatment. CLPP knockdown in MEL-20- 06-039 resulted in reduced sensitivity to growth inhibition by ONC212, supporting that the inhibition of cell viability by ONC212 was mediated via CLPP activation (Figure S2).

ONC212 also inhibited migration of UM cell lines, albeit to different degrees (Figures 3A and S3). Moreover, ONC212 treatment reduced the expression of cell migration markers β1-integrin and F-actin, as shown by western blotting and quantitation of their expression (Figure 3B).

**Figure 3.**
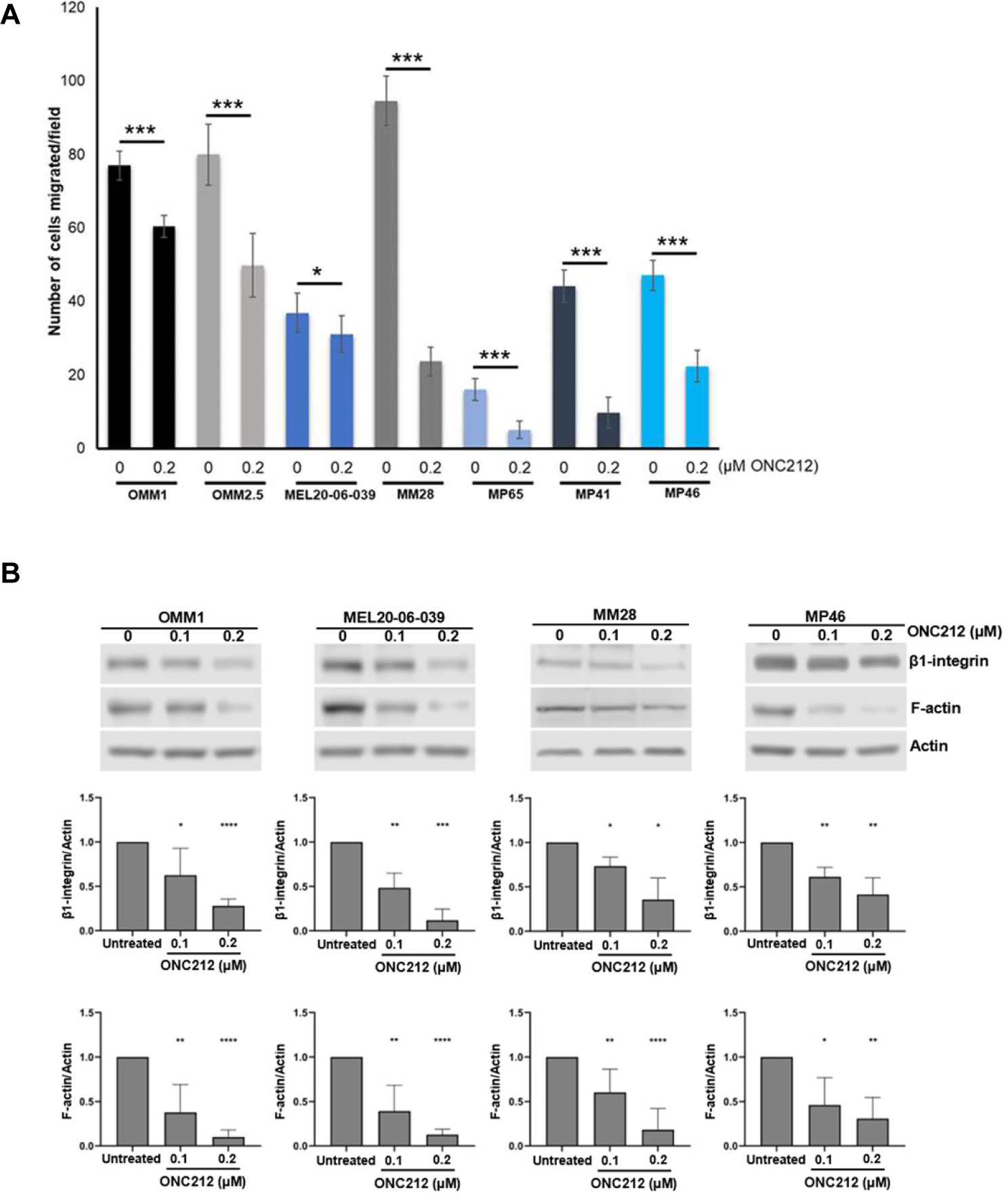
ONC212 inhibits UM cell migration. (A) In vitro cell migration assay with ONC212 treatment (0.2 µM) of UM cell lines (OMM1, OMM2.5, MEL20-06-039, MM28, MP65, MP41, and MP46). An average of five fields of cells/filter were counted under a microscope with 40X magnification. The average number of cells counted/field were obtained in two independent experiments and the mean ± SD of these average cell counts/field were plotted as bar graphs. p- values were calculated by comparing untreated vs. ONC212 treated cells (* p < 0.05; ** p < 0.01; *** p < 0.001). **(B)** Western blots showing the changes in cell migration markers, β1-integrin, and F-actin upon ONC212 treatment in OMM1, MEL20-06-039, MP46 and MM28 cell lines (top panels), and corresponding quantitation of these proteins normalized with β-actin loading control (bottom panels; * p < 0.05; ** p < 0.01; *** p < 0.001).

### ONC212 treatment impacts major metabolic and cell survival pathways in UM

As OXPHOS is one of the primary energy metabolism pathways in UM cells, and a key regulator of cell survival and therapeutic resistance, we evaluated the effect of ONC212 broadly on protein signaling pathways. UM cell lines (MEL20-06-039, MP41, MEL270, MP46, OMM2.3, and OMM2.5) were treated with 0.1 and 0.2 µM ONC212 for 48 h. RPPA analysis identified significant alterations in several important cell signaling proteins in all the cell lines tested (Figure 4A and S4A). For clarity and space economy, representative data from two cell lines (MEL-20- 06-039 and OMM2.3) are shown in Figure 4A. The normalized linear data shows a significant dose-dependent decrease in SDHA and SDHB in all the cell lines tested (p<0.05), which indicates that ONC212 impacts OXPHOS effector levels as anticipated and is consistent with our western blotting analysis (Figure 2A). ONC212 treatment increased levels of p85, the inhibitory subunit of PI3K catalytic activity. ONC212 also increased AMPK and decreased mTOR activation in all UM cell lines tested (p<0.05), suggesting ONC212 induces metabolic stress. Glucose metabolism modulators such as DUSP6 and AKT2 were also decreased with ONC212 treatment (p<0.05). Retinoblastoma (Rb) protein phosphorylation at S807-811 was decreased (p<0.05). This Rb phosphorylation regulates interaction of Rb with pro-apoptotic BAX protein (40). These results suggest that ONC212 treatment affects major metabolic and apoptotic proteins in UM cells.

**Figure 4.**
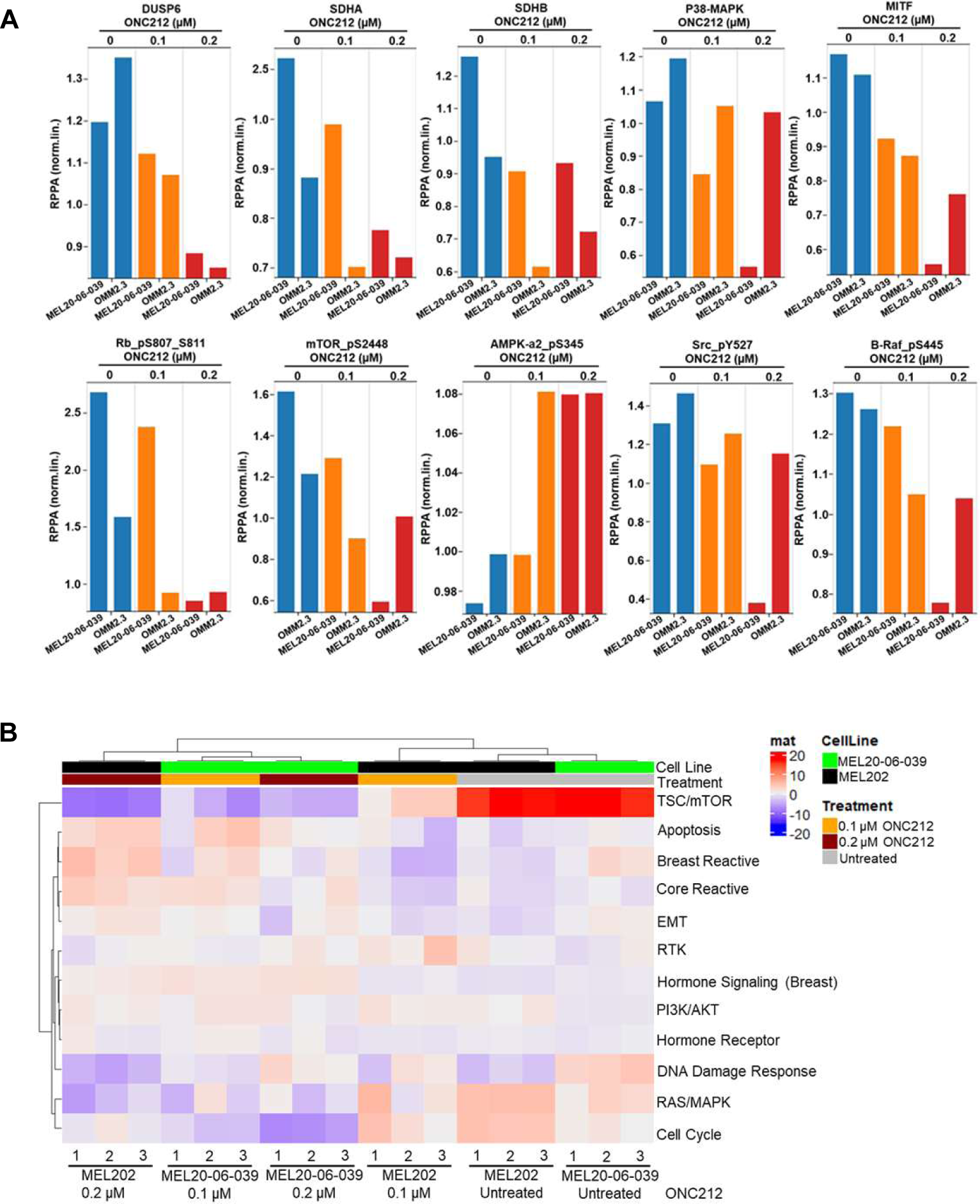
Alterations in protein profiles of UM cells with ONC212 treatment. (A) The top ten altered proteins from RPPA profiling of MEL-20-06-039 and OMM2.3 UM cell lines. The RPPA expression values are displayed on the y-axis for antibodies that show a significant difference (p < 0.05) compared to control sample. Cells were treated either 48 h with 0.1 or 0.2 µM concentration of ONC212, and untreated cells are used as controls. **(B)** A heatmap of cellular signaling pathway activity scores after 48 h of ONC212 vs. vehicle treatment of UM cell lines (MEL20-06-039 and MEL202).

We performed pathway analysis on RPPA data for MEL202 (D3) and MEL20-06-039 (M3) UM cell lines after treatment with ONC212 (0.1 and 0.2 µM) in triplicate for 24 and 48 h (Figure 4B and S4B). ONC212 treatment downregulated mTOR signaling (≥20-fold) and MAPK pathway signaling (≥10-fold) and upregulated apoptotic pathways (∼10-fold). Additionally, we observed a significant difference in the expression of proteins involved in the DNA damage response between the D3 and M3 cell lines at baseline. Therefore, the RPPA data indicates that ONC212 impacts UM cellular metabolism and survival.

### ONC212 inhibits protein metabolism and lipid biosynthesis in UM

Our prior work showed that M3UM relies on high OXPHOS and high expression of mitochondrial OXPHOS effectors for survival (16). Since ONC212 treatment inhibited UM cell survival (Figure 2 ), we investigated whether this treatment alters the metabolism of UM cells. MEL20-06-039 cells were treated with ONC212 (0.1 and 0.2 µM), and the metabolites were extracted for high-resolution mass spectrometry-based global metabolomics and lipidomics profiling. Global pathway analysis showed that ONC212 treatment significantly reduces energy metabolism pathways in a dose-dependent manner (Figure 5A and S5A). ONC212 treatment diminished ROS regulating metabolites involved in glutathione, aspartate-glutamate, and NADH metabolism (Figure 5A). ONC212 also reduced the levels of the essential amino acids- with the exception of an increase in levels of Valine, Lysine, Tyrosine, and Arginine (Figure 5B), which may be involved as intermediates in perturbation of protein metabolism upon ONC212 treatment. Metabolites involved in redox metabolism were also markedly reduced with ONC212 treatment (Figure 5C), suggesting that ONC212 affects amino acid and redox metabolism. Moreover, ONC212 decreased metabolites involved in macromolecule metabolism such as nucleotide biosynthesis and amino sugar metabolism (Figure S5B). In contrast, ONC212 treatment resulted in enrichment of a set of signature metabolites including itaconic acid, lactic acid, and TCA cycle intermediates (Figure S5C). These metabolites and their downstream networks are often involved in cell death and inflammation in response to drug treatment(41).

**Figure 5.**
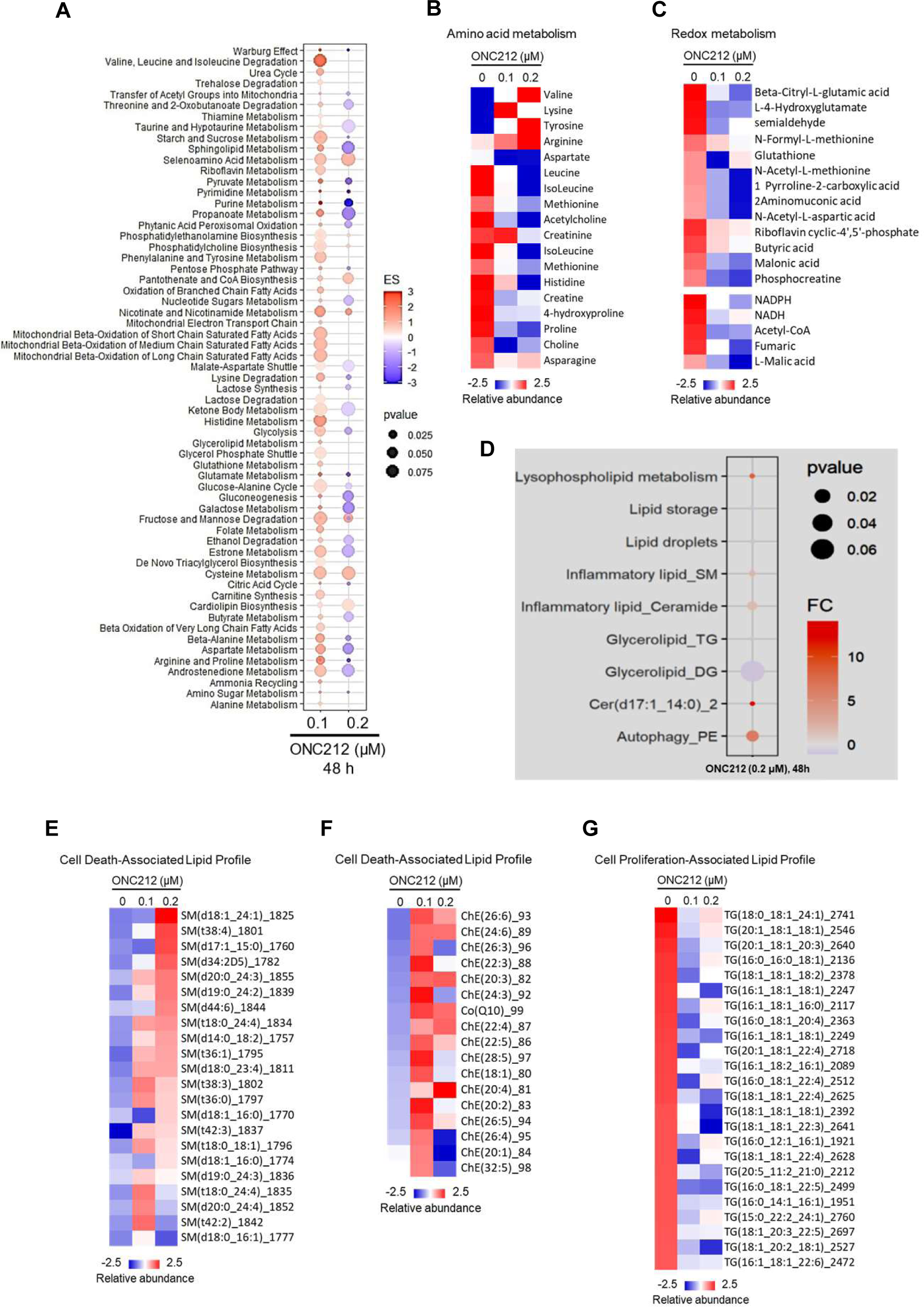
Alterations in metabolomic and lipidomic profiles of UM cells with ONC212 treatment. (A-G) Mass spectrometry-based global metabolomics and lipidomics of UM cell line MEL20-06-039, treated with 0.1 and 0.2 µM of ONC212 for 48 h. **(A)** Pathway analysis and trends from significant changes observed in the metabolic profile. **(B-C)** Heatmap showing changes in essential amino acids **(B)**, and changes in metabolites involved in redox metabolism **(C**). **(D)** Pathway analysis and trends using lipidomics profiling showing significant changes upon ONC212 treatment. **(E-G)** Heatmaps of lipid profiles analysis in Sphingomyelin (SM) **(E)**, in inflammatory lipid signature such as ceramide, cholesterol ester, and CoQ **(F)**, and in triglyceride signature (TG) **(G)**.

Pathway enrichment analysis revealed differential enrichment of lipid ontologies by ONC212 (Figure 5D). ONC212 increased levels of cell-death and inflammatory lipids like sphingomyelin (SM) (Figure 5E), cholesterol ester (ChE), and coenzyme Q (CoQ) (Figure 5F). In contrast, lipids such as triglycerides (TG), which is essential for cell proliferation, decreased (Figure 5G). The detailed heatmaps of the global lipidomics profiling are provided in Figure S5D-F. Collectively, ONC212 alters the UM cell lipidome by elevating lipids linked to inflammation and cell death while suppressing proliferative lipids(42).

### ONC212 inhibits UM liver metastasis growth and improves survival *in vivo*

To study the effect of ONC212 in a clinically relevant model, we standardized an orthotopic mouse model of UM liver metastasis. Luciferase tagged UM cells (92.1, MEL20-06-039, MP41 and MM28) were injected into the spleen of NSG mice, followed by splenectomy (37). After confirming tumor growth in the liver, mice were randomized and treated with 25 mg/kg ONC212 or vehicle twice weekly. Tumor burden was monitored through weekly IVIS scans (Figures 6A and S6A).

**Figure 6.**
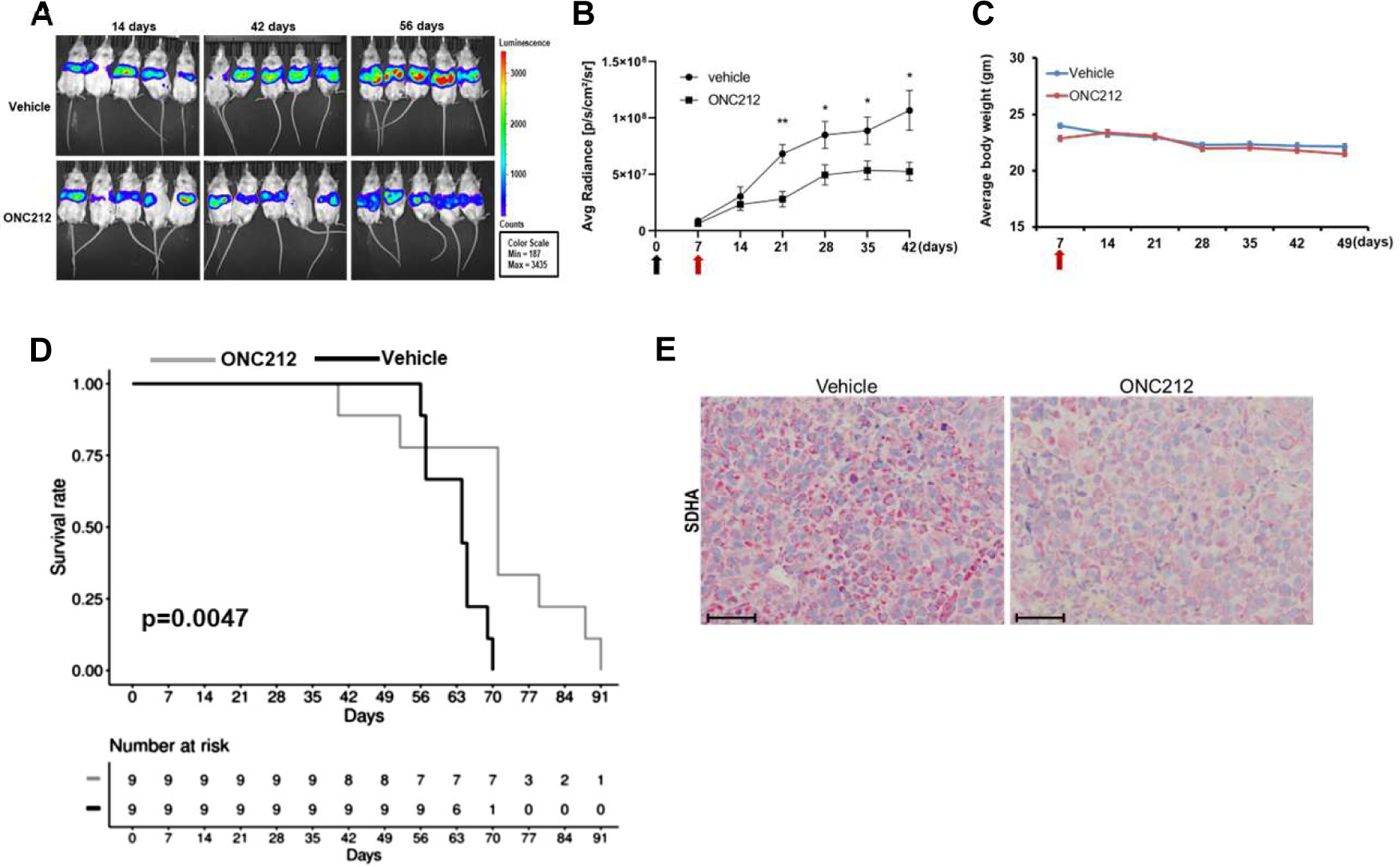
ONC212 reduces tumor burden and improves survival in UM liver metastasis model. (A) Representative images of weekly bioluminescence scans of mice with liver metastases of UM 92.1 treated with vehicle (control) or ONC212 (25 mg/kg). **(B)** Tumor growth curves plotted from bioluminescence scans post-tumor initiation. The black arrow represents the splenic injection and red arrow indicates beginning of treatment. **(C)** Mouse body weight with vehicle and ONC212 treatment; red arrow indicates beginning of treatment. **(D)** Kaplan-Meier plots of 92.1 model with ONC212 treatment compared to vehicle treated controls; n = 10 per treatment group; 92.1 vehicle vs. ONC212 p = 0.0047; Hazard ratio from Cox proportional hazards model: 7.80 [95% confidence interval: 1.573 to 38.66]. **(E)** Representative images (scale bar = 100 µm) of SDHA immunohistochemical staining in vehicle and ONC212-treated 92.1 tumors in liver.

Treatment with ONC212 significantly reduced liver tumor burden of hepatic xenografts of 92.1 and MEL20-06-039 (Figures 6B and S6B) and improved overall survival (Figures 6D and S6D). Of note, with prolonged ONC212 treatment (9-10 weeks) we observed tumor recurrence, indicating the development of ONC212 resistance (Figures 6A and S6A). No significant changes in mouse body weight were observed during ONC212 treatment (Figure 6C and S6C). ONC212 did not improve survival in mice with hepatic xenografts of MP41 and MM28 (Figure S7). Mouse livers were excised at the end of the survival analysis for further study. IHC analysis showed a reduction in SDHA protein in the ONC212-treated vs. vehicle treated tumors (Figure 6E and S6E). Taken together, our data indicates a beneficial treatment effect of ONC212 in 2 of 4 models of UM liver metastasis.

## Discussion

There is a critical need for new effective strategies for UM. Previous studies by our group, subsequently and independently validated by others, identified elevated mitochondrial OXPHOS and high expression of OXPHOS effector proteins in mUM (16,17,43). Upon targeting OXPHOS directly using IACS10759 in UM cell lines, we observed limited reduction of cell viability. This poor response was due to elevated expression of OXPHOS effectors. Moreover, a phase 1 clinical study with IACS10759 resulted in unacceptable toxicities (NCT 03291938) (19). Therefore, there was a necessity for a safe and effective alternate strategy to target the OXPHOS-dependency of UM. In this study, we show that the imipridones ONC201 and ONC212, which inhibit OXPHOS indirectly by activating mitochondrial CLPP, reduce OXPHOS effectors; inhibit UM cell survival and migration; and induce cell death *in vitro.* ONC212 treatment also affected multiple signaling and metabolic pathways in UM cells. Further, ONC212 significantly reduced tumor burden and improved survival in two of four *in vivo* models of UM liver metastasis. Together these results support that imipridones may be an important and a novel strategy for mUM.

This is the first attempt, to our knowledge, to target UM growth using imipridones. Though CLPP has been identified as the target of ONC212, the mechanism of cancer cell death induced by ONC212 is not well-understood (44). To identify mechanism(s) of ONC212 action in UM cells, we analyzed multiomics data, including proteomic, metabolomic and lipidomic profiles. These analyses highlighted the activation of apoptotic pathways and induction of metabolic stress by ONC212 treatment. In pancreatic cancer, ONC212 has been shown to collapse mitochondrial function (44). We observed that the mitochondrial proteins SDHA and SDHB were significantly reduced by ONC212, consistent with CLPP activation. Additionally, we observed alterations to AMPK and mTOR levels. AMPK and mTOR are master regulators of cellular metabolism, as they sense and respond to metabolic environment and oxidative stress. Activated AMPK has also been shown to inhibit mTORC1 signaling (45). Our observation of higher AMPK phosphorylation and lower mTOR with ONC212 treatment suggests an induction of metabolic stress in these cells.

Since our proteomics data suggests that ONC212 reduces the levels of OXPHOS effectors and induces metabolic stress in UM cells, we further dissected the metabolic effects of ONC212 treatment in UM cells. Global metabolomics and lipidomics analysis identified downregulation of ROS neutralizing entities such as glutathione and polyamines by ONC212, implying ROS upregulation is involved in cell death with this treatment. There was also a significant impairment of protein biosynthesis, which may further contribute to ONC212-induced cell death. Our metabolic profile analysis also shows a reduction in Warburg effect. An investigation of the lipid profiles show an accumulation of inflammatory lipids that are known to induce cell death (46,47). Moreover, a reduction in triglycerides signify an inhibition of the cell proliferation pathways with ONC212 treatment.

Perhaps most importantly, our studies show that ONC212 reduced UM tumor growth *in vivo* in two different orthotopic mouse models for mUM liver metastasis. The liver is the most frequent site of UM metastasis, and liver involvement has been associated with poor outcomes with other therapies, including immune checkpoint inhibitors. Thus, the observed results with single-agent ONC212 in this aggressive, clinically relevant model are promising and clinically relevant. However, there is clearly a need to build upon this initial demonstration. The heterogeneous outcomes observed in the 4 UM models tested, along with the eventual development of resistance in the models that demonstrated initial benefit, provide a foundation for further characterization of the determinants of the efficacy of ONC212 for mUM- and hopefully the development of even more effective combinatorial strategies.

## Acknowledgments

We thank The University of Texas MD Anderson Flow Cytometry and Cellular Imaging Facility supported in part by NIH (CA016672); The Metabolomics Facility supported in part by The University of Texas MD Anderson Cancer Center and P30CA016672; and The University of Texas MD Anderson RPPA Core, supported by NCI Grant # CA16672 and NIH grant R50CA221675. The patient tissues for IHC were provided by MelCore database supported by The University of Texas MD Anderson Melanoma SPORE (P50CA221703) Chimerix/Oncoceutics provided ONC201 and ONC212 for these studies.

## Author Contributions

Concept and design: C.C.; Acquisition of data: C.C., R.B., F.S.K., M.M.A., M.A., R.R.M., R.L.S.; Analysis and interpretation of data: C.C., J.R., I.M., M.M.A., M.A., S.Y.; Writing, review, and/or revision of the manuscript: C.C., J.R., S.P.P., R.B. I.M., S.Y., M.A.D.; Manuscript review: E.A.G.; Study supervision: C.C. All authors have read and agreed to the published version of the manuscript.

## Translational Relevance

There remains an unmet need for new, more effective treatments for metastatic uveal melanoma (UM). We recently showed that UM is characterized by a high OXPHOS metabolic phenotype. Here we have shown that the imipridones ONC201 and ONC212, which are small molecule activators of mitochondrial CLPP and have demonstrated safety in cancer patients, are able to inhibit OXHOS effectors and UM cell growth *in vitro*. Further, ONC212 inhibited UM liver metastases *in vivo*. These promising findings suggest that imipridones may be a novel and actionable therapeutic strategy for UM.

## Supplementary Data Figure Legends

**Supplementary Figure 1.**
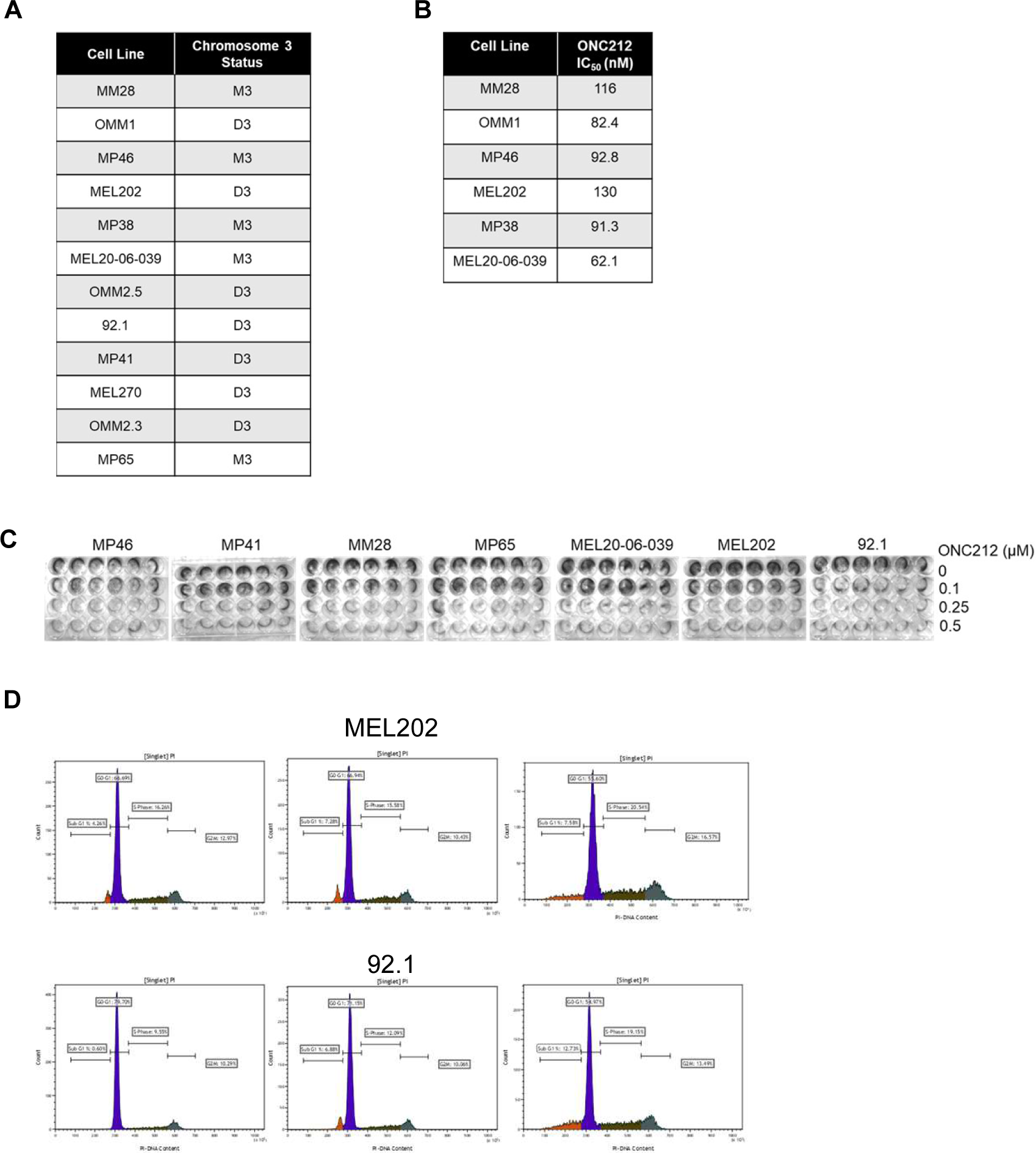
UM cell lines used in this study and the effect of ONC212 on UM cell survival (A) Chromosome 3 status of UM cell lines used in this study. (**B**) IC_50_ values of ONC212 in UM cell lines calculated from MTT assays. (**C**) Colony formation assay: Representative images of crystal violet staining shows fewer colonies formed by UM cell lines (MEL20-06-039, 92.1, MP46, MEL202, MM28, MP65, and MP41) with ONC212 treatment (0.1, 0.25, and 0.5 µM for seven days) compared to untreated controls. **(D)** Representative FACS histograms of DNA content detected by propidium iodide staining to detect cell cycle status in UM cell lines (MEL202 and 92.1) show a dose-dependent increase in the number of cells in G0/G1 phase with ONC212 treatment (0.1 and 0.2 µM for 72 h) vs. no treatment.

**Supplementary Figure 2.**
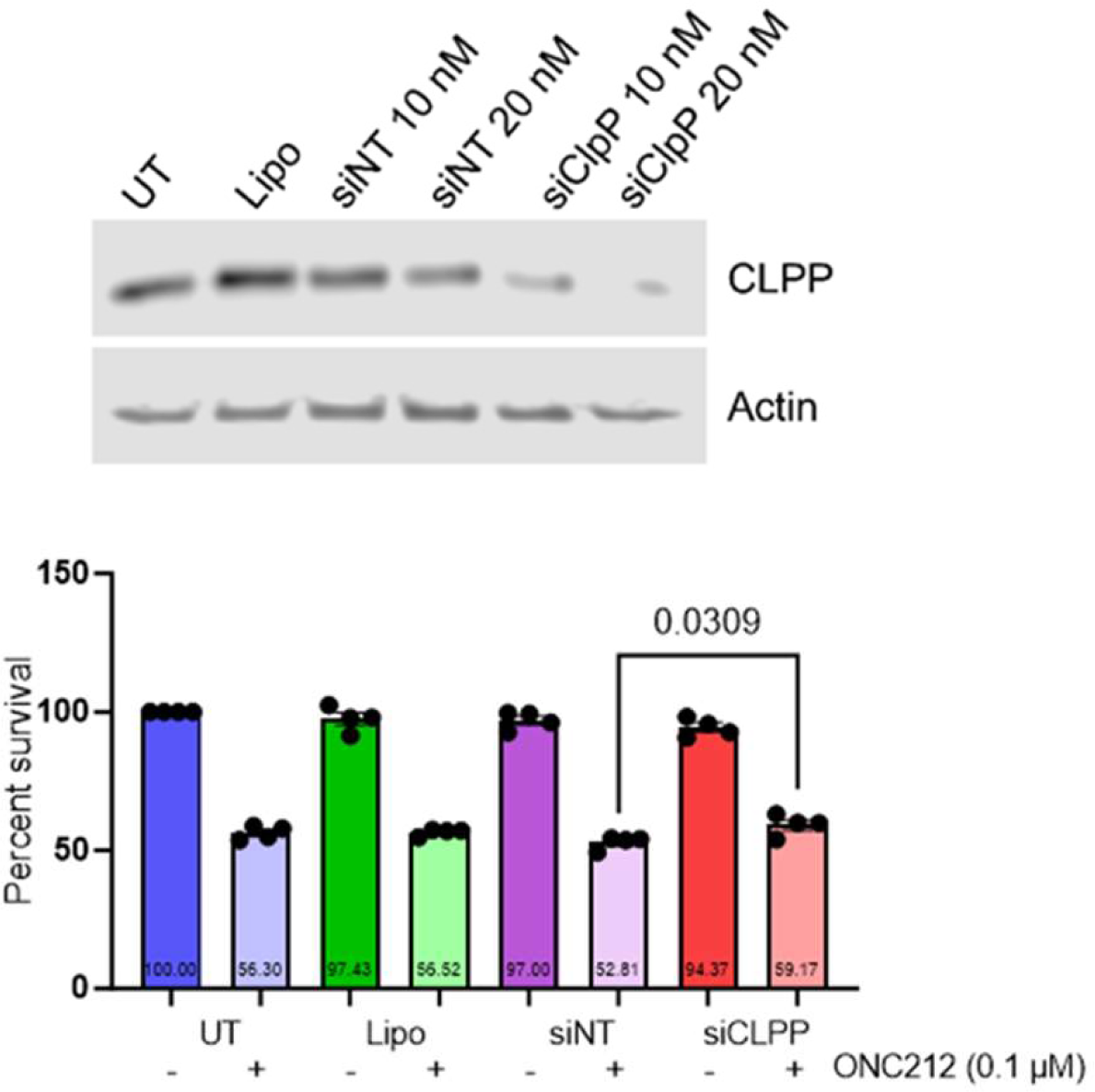
CLPP as ONC212 target in UM cells. (A) ClpP knockdown using siRNA (10 and 20 nM siClpP) and western blot analysis showing CLPP expression levels. Actin expression served as loading control. **(B)** Cell viability assay post-ClpP knockdown (10 nM) and 0.1 µM ONC212 treatment in MEL20-06-039 cells. The controls were Untreated (UT), Lipofectamine-treated (Lipo), Non-targeting siRNA-treated (siNT) cells. p = 0.0309 for comparison between siNT and siClpP with ONC212 treatment.

**Supplementary Figure 3.**
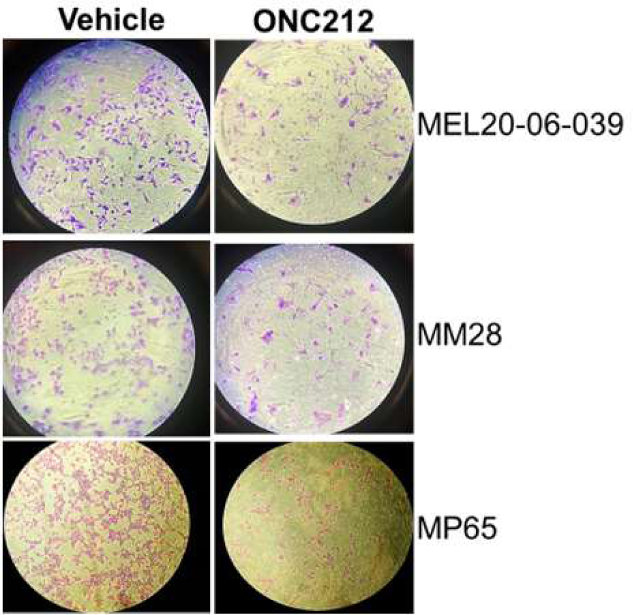
ONC212 inhibits UM cell migration. Representative brightfield images (10X) of cell migration assay in UM cell lines, MEL20-06-039, MM28, and MP65, treated with vehicle or 0.2 µM ONC212.

**Supplementary Figure 4.**
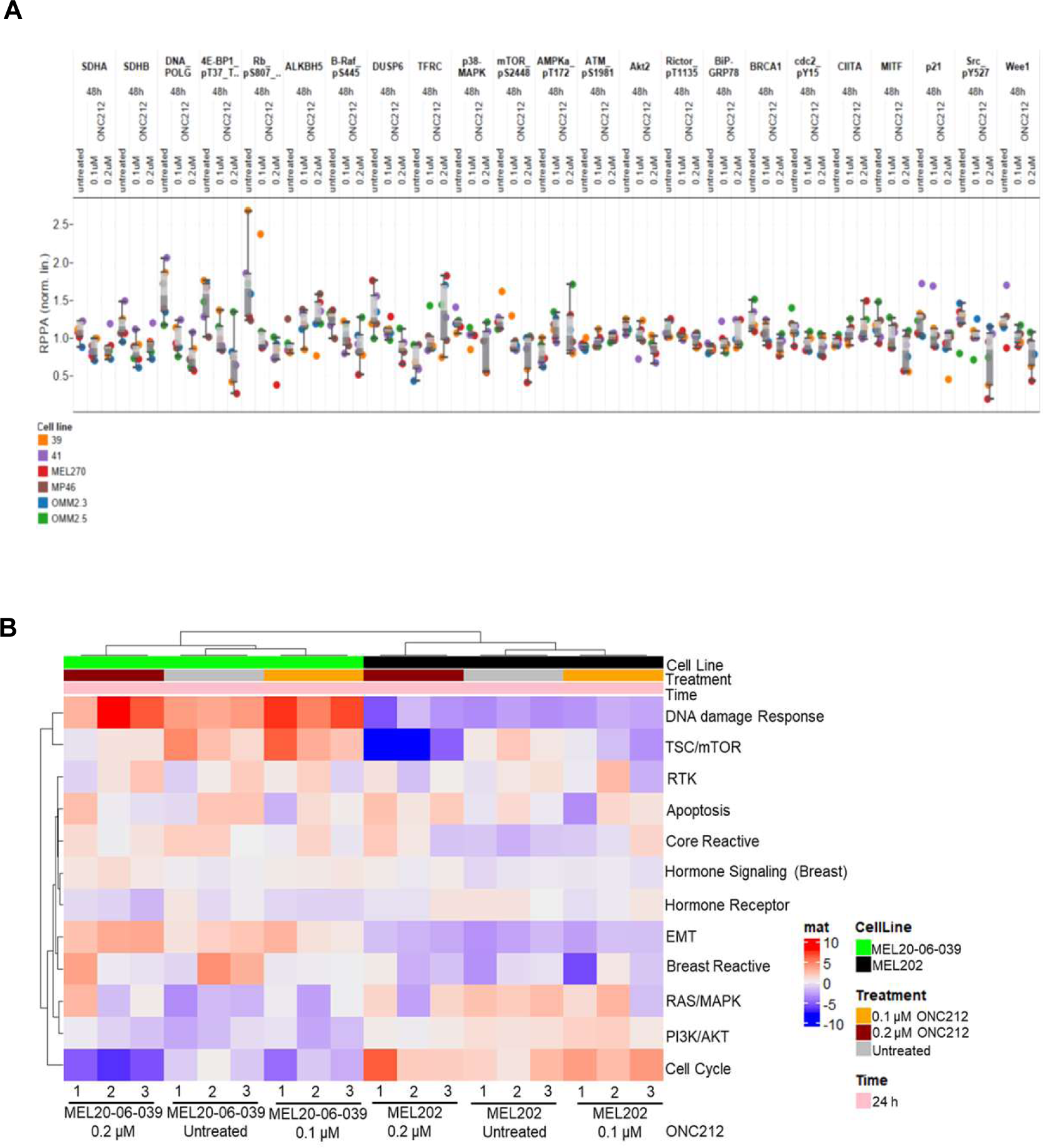
ONC212 alters global proteomics of UM cells. (A) RPPA protein profiling of four UM cell lines (MEL-20-06-039, MP41, MEL270, MP46, OMM2.3, and OMM2.5, represented by heatmaps. Cells were treated for 48 h with either 0.1 or 0.2 µM ONC212. **(B)** 24 h RPPA data set supplementing Figure 4B.

**Supplementary Figure 5.**
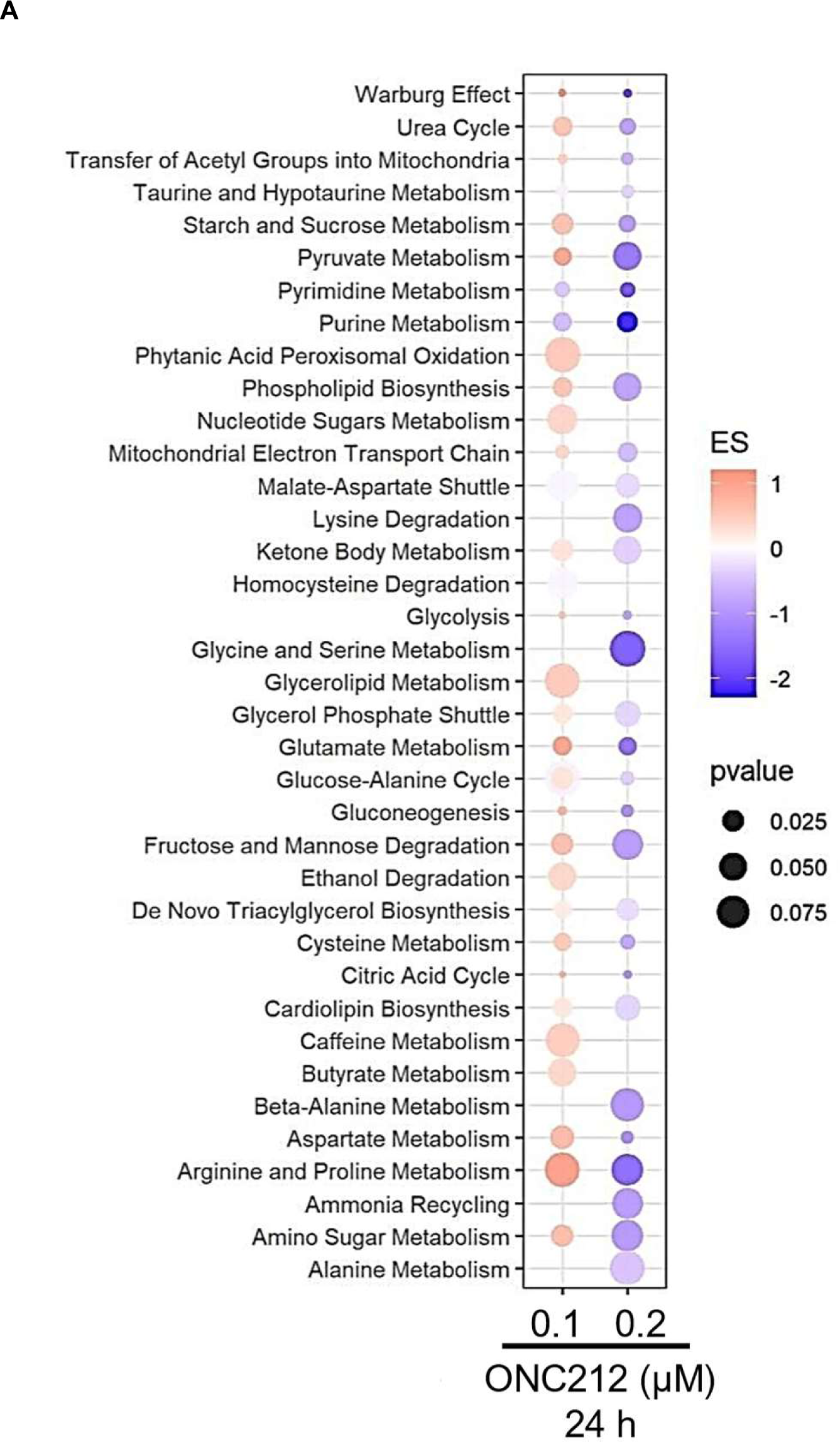
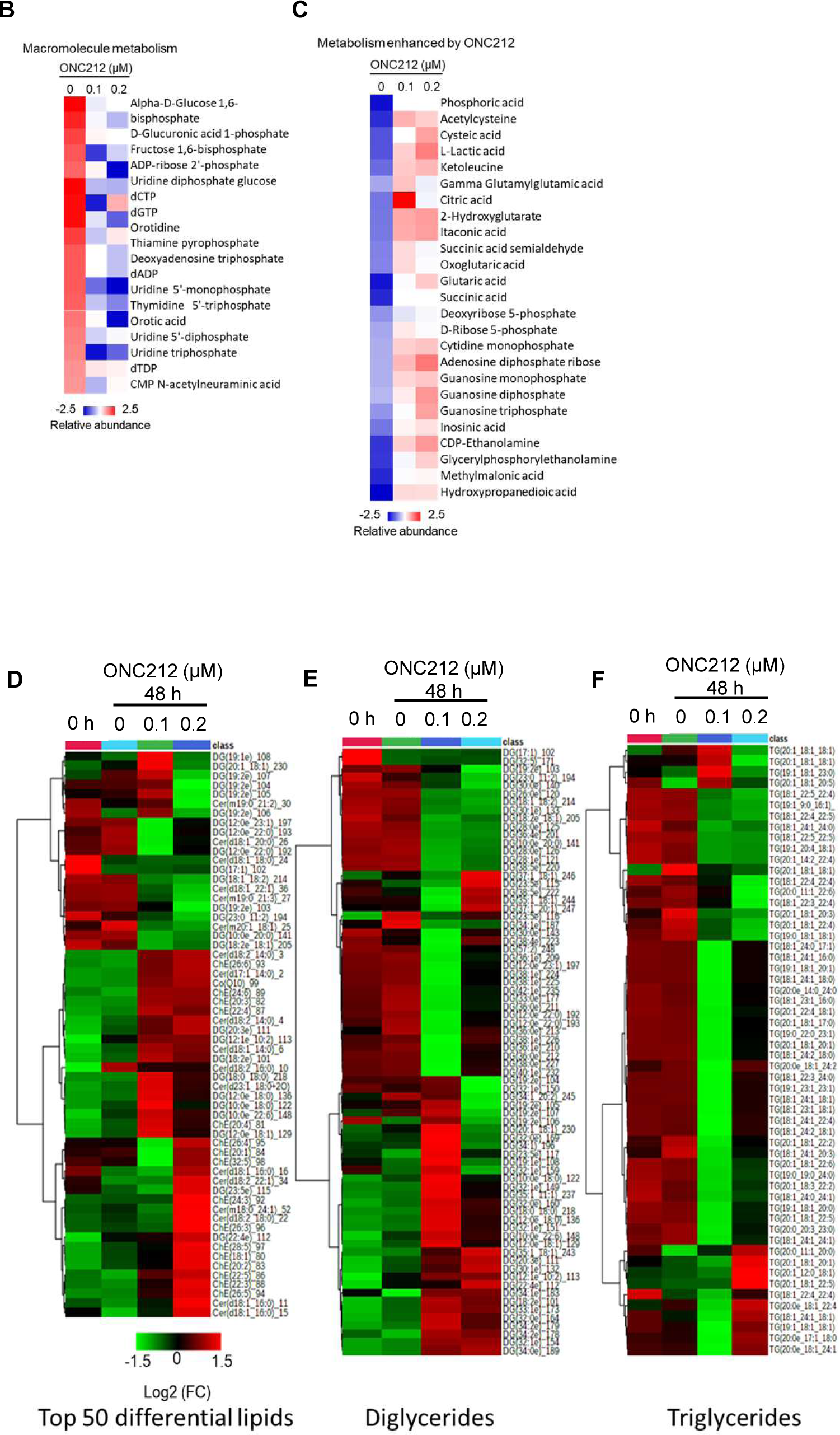
ONC212 alters global metabolomic and lipidomics profiles of UM cells. Global metabolomics and lipidomics profiling of MEL20-06-039 cells treated with 0.1 and 0.2 µM of ONC212 for 24 h. **(A)** Pathway analysis and trends from significant changes observed in the metabolic profile shows the relative utilization of individual metabolic pathways. **(B and C)** Heatmaps of metabolites related to macromolecule metabolism **(B)** and metabolites enhanced by ONC212 treatment **(C)**, obtained from mass spectrometry-based global metabolomics **(D-F)** Heatmaps of lipid profiles showing normalized abundance of lipids, di and triglycerides.

**Supplementary Figure 6.**
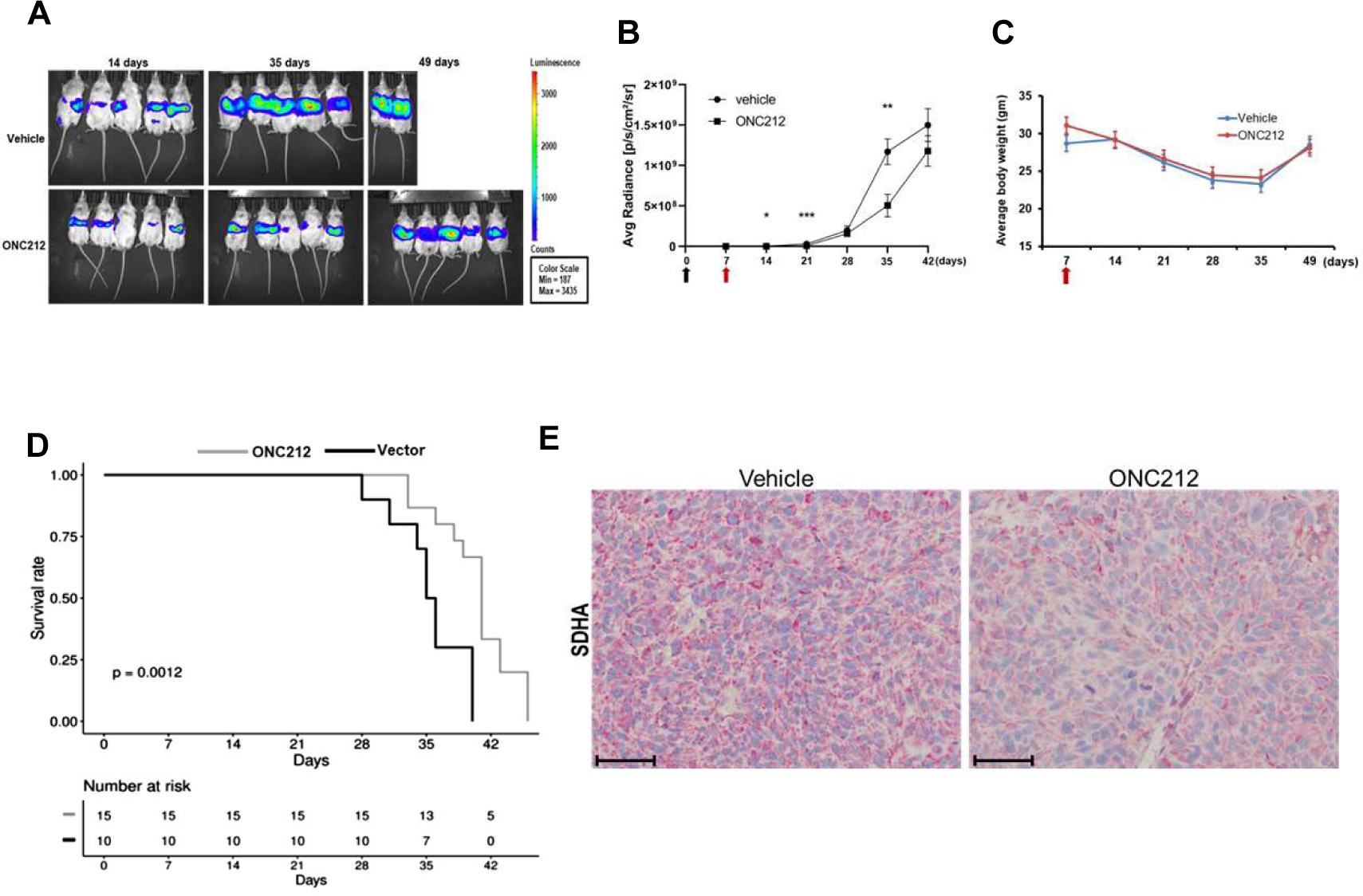
ONC212 reduces tumor burden and improves survival in MEL20-06-039 orthotopic liver metastases model. (A) Representative images of weekly bioluminescence scans of MEL20-06- 039 liver metastasis mouse models treated with vehicle (control) and ONC212 (25 mg/kg). **(B)** Tumor growth curves plotted from bioluminescence scans post-tumor initiation. The black arrow represents the splenic injection and red arrow indicates beginning of treatment. **(C)** A plot of mouse body weight in vehicle and treatment groups over the period of experiment; red arrow indicates beginning of treatment. **(D)** Kaplan-Meier plots of MEL20-06-039 model after ONC212 treatment compared to vehicle treated controls; n = 10 per treatment group; MEL20-06-039 vehicle vs. ONC212 p = 0.0012; Hazard ratio from Cox proportional hazards model: 10.60 [95% confidence interval: 2.205 to 50.96] **(E)** Representative images (scale bar = 100 µm) of SDHA immunohistochemical staining in vehicle and ONC212-treated MEL20-06-039 tumors in liver.

**Supplementary Figure 7.**
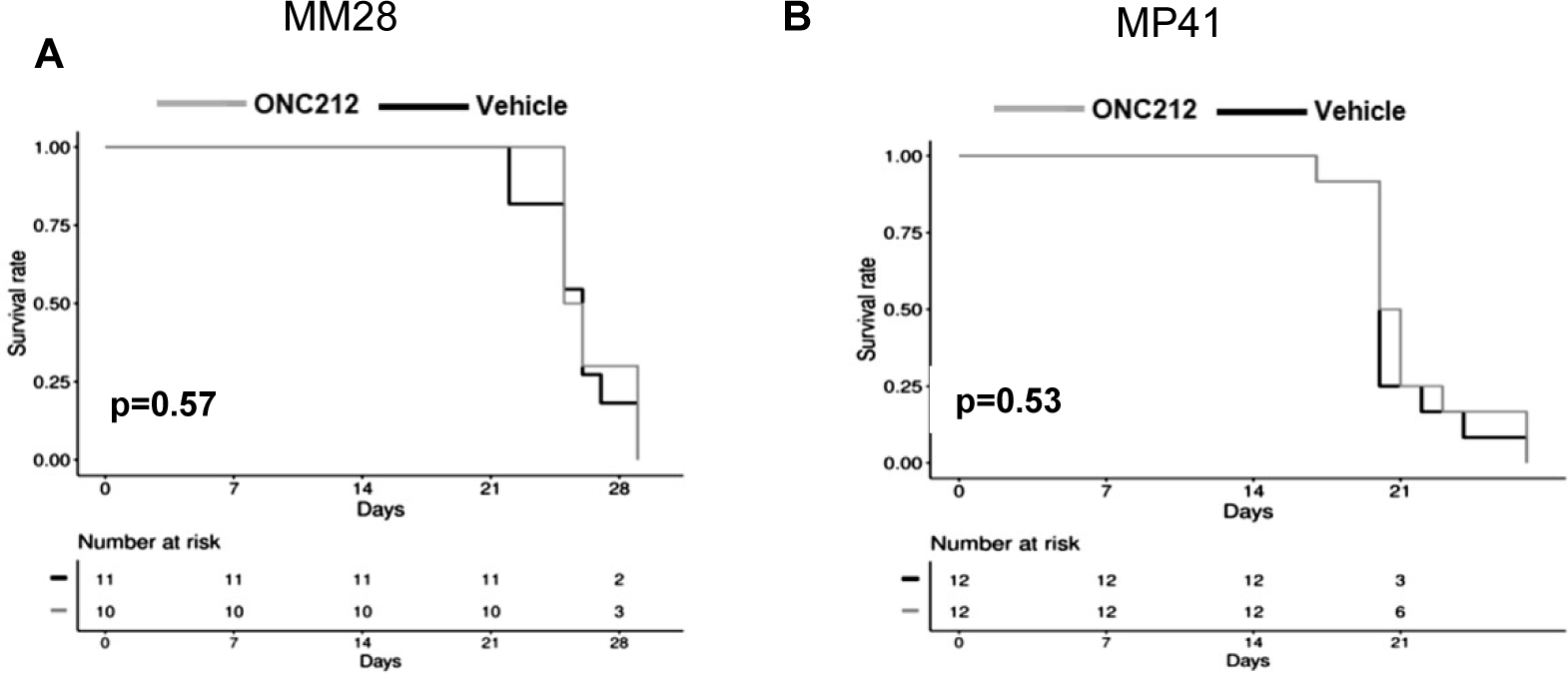
ONC212 does not improve survival in MM28 and MP46 orthotopic liver metastases models. Kaplan-Meier plots and corresponding Hazard Ratios of **(A)** MM28 and **(B)** MP41 models with ONC212 treatment (25 mg/kg, twice weekly) compared to vehicle treated controls; n = 10 per treatment group. The p-values for MM28 model and MP41 model in vehicle vs. ONC212 treatment are p = 0.57 and p = 0.53, respectively.

